# PKA-mediated phosphorylation of Neuroligin-2 regulates its cell surface expression and synaptic stabilisation

**DOI:** 10.1101/2020.07.23.218008

**Authors:** Els F. Halff, Saad Hannan, Trevor G. Smart, Josef T. Kittler

## Abstract

The trans-synaptic adhesion molecule Neuroligin-2 (NL2) is essential for the development and function of inhibitory synapses. NL2 recruits the postsynaptic scaffold protein gephyrin, which in turn stabilises GABA_A_ receptors (GABA_A_Rs) in the postsynaptic domain. Dynamic regulation of synaptic GABA_A_R concentration is crucial for inhibitory neurotransmission efficacy. Changes in synaptic levels of NL2 contribute to regulating GABA_A_R synaptic concentration, however the mechanisms that control NL2 synaptic stabilisation are mostly unknown. Here, by combining biochemistry, imaging, single particle tracking and electrophysiology, we identify a key role for cAMP-dependent protein kinase (PKA) in synaptic stabilisation of NL2. We show that PKA-mediated phosphorylation of NL2 at S714 causes its dispersal from the synapse and reduces NL2 surface levels, leading to a loss of synaptic GABA_A_Rs. Conversely, enhanced stability of NL2 at synapses through abolishing phosphorylation leads to increased inhibitory signalling. Thus, PKA plays a key role in regulating NL2 function and synaptic inhibition.

## Introduction

Synaptic inhibition through postsynaptic γ-aminobutyric acid (GABA) type-A receptors (GABA_A_Rs) is crucial for controlling the balance between excitation and inhibition (E/I) of neuronal signaling in the brain. Defects in neuronal inhibition underlie several neurological conditions including epilepsy, autism and schizophrenia (Smith and Kittler, 2010, Bozzi et al., 2018, Charych et al., 2009). To maintain the E/I balance during neural activity, the strength of the postsynaptic response is dynamically regulated through altering the number of GABA_A_Rs; this involves processes such as phosphorylation and ubiquitination, which impact on stabilisation *versus* dispersal of receptors from the synapse and receptor internalisation *versus* recycling (Luscher et al., 2011, Arancibia-Carcamo and Kittler, 2009, Vithlani and Moss, 2009, Kittler et al., 2005, Nakamura et al., 2015, Muir et al., 2010). GABA_A_R function is furthermore regulated by the auxiliary protein Shisa7 (Han et al., 2019).

The postsynaptic membrane-spanning adhesion molecule Neuroligin-2 (NL2), together with its postsynaptic binding partners LHFPL4 (Lipoma HMGIC Fusion Partner-Like 4) and Slitrk3 (Slit- and Trk-like family protein 3), plays a key role in the development and function of GABAergic synapses (Davenport et al., 2017, Wu et al., 2018, Li et al., 2017, Sudhof, 2018, Yamasaki et al., 2017). Crucially, NL2, through direct interaction with collybistin, drives clustering of a post-synaptic gephyrin scaffold, which stabilises GABA_A_Rs at synapses (Poulopoulos et al., 2009, Tyagarajan and Fritschy, 2014). Furthermore, NL2 ensures correct positioning opposite pre-synaptic terminals through interaction with pre-synaptic neurexins (Bemben et al., 2015, Varoqueaux et al., 2004, Craig and Kang, 2007).

Although changes in synaptic levels of NL2 contribute to the modulation of inhibitory synapse size and signalling strength (Halff et al., 2019, Binda et al., 2019, Chubykin et al., 2007, Bemben et al., 2015), we do not currently understand what molecular mechanisms regulate synaptic stabilisation of NL2 as well as its internalisation. By comparison, Neuroligin-1 (NL1) has two distinct phosphorylation sites, regulated by CamKII and PKA respectively, which regulate its surface levels and modulate excitatory signalling (Bemben et al., 2014, Jeong et al., 2019). In NL2, phosphorylation at S714 induces recruitment of the peptidyl-prolyl *cis-trans* isomerase protein Pin1. Downstream of phosphorylation, Pin1 activity disrupts interaction between NL2 and gephyrin, whereas knockout of Pin1 leads to increased inhibitory synapse size and enhanced GABAergic signalling (Antonelli et al., 2014). Together, these studies point towards a role for NL2 phosphorylation in regulating inhibitory signalling.

Whereas these studies investigated downstream consequences of NL2 phosphorylation, it is still unknown what pathway leads to phosphorylation of NL2, as well as the consequences of phosphorylation for localisation and synaptic stability of NL2 itself. In this study we investigated the molecular mechanisms inducing NL2 phosphorylation, and its effect on NL2 dynamic behaviour. We show that S714 is a unique serine phosphorylation site that is constitutively phosphorylated in neurons. Using *in vivo* and *in vitro* methods, we identify cAMP-dependent protein kinase (PKA) as the kinase phosphorylating NL2 at S714. Moreover, we show that PKA-induced phosphorylation of NL2 causes its rapid dispersal from the synapse and reduces its surface levels, consequently reducing synaptic GABA_A_R levels. Conversely, mutation of the phosphorylation site leads to reduced internalisation and enhances inhibitory signalling. Thus, our study establishes a role for PKA-mediated phosphorylation in the surface and synaptic stabilisation of NL2 and modulation of inhibitory signalling.

## Results

### NL2 is uniquely and constitutively phosphorylated at S714

Mass spectrometry studies aiming to map tissue-specific phospho-proteomes identified residues S714, S719 and S721 in the intracellular region of murine and rat NL2 as putative serine phosphorylation sites (Huttlin et al., 2010, Li et al., 2016, Lundby et al., 2012). To investigate the extent and role of phosphorylation at these sites, we created phospho-null (S to A) and phospho-mimetic (S to D) mutants of murine HA-tagged NL2 (^HA^NL2; Fig.1a). As a control we also included a non-related mutant, Y770A; this tyrosine is conserved across Neuroligins, and phosphorylation of the equivalent site in NL1 (Y782) has been shown to prevent gephyrin-binding (Giannone et al., 2013). We expressed WT and mutant ^HA^NL2 in COS cells and analysed cell lysates by western blotting, using a standard SDS-PAGE gel supplemented with PhosTag. This alkoxide-bridged Mn^2+^-binding compound, when added to a standard polyacrylamide gel, reduces the migration of phosphorylated proteins compared to their non-phosphorylated counterparts (Kinoshita et al., 2006), thus enabling detection of protein phosphorylation without the requirement of generating site-specific phospho-antibodies. Using this method, we found that WT and Y770A ^HA^NL2 are phosphorylated at steady state, whereas mutants of S714 alone or in combination with S719 and S721 are not phosphorylated (Fig.1b). This indicates that ^HA^NL2 is basally phosphorylated at S714, while the other sites are not phosphorylated in COS cells.

**Figure 1:**
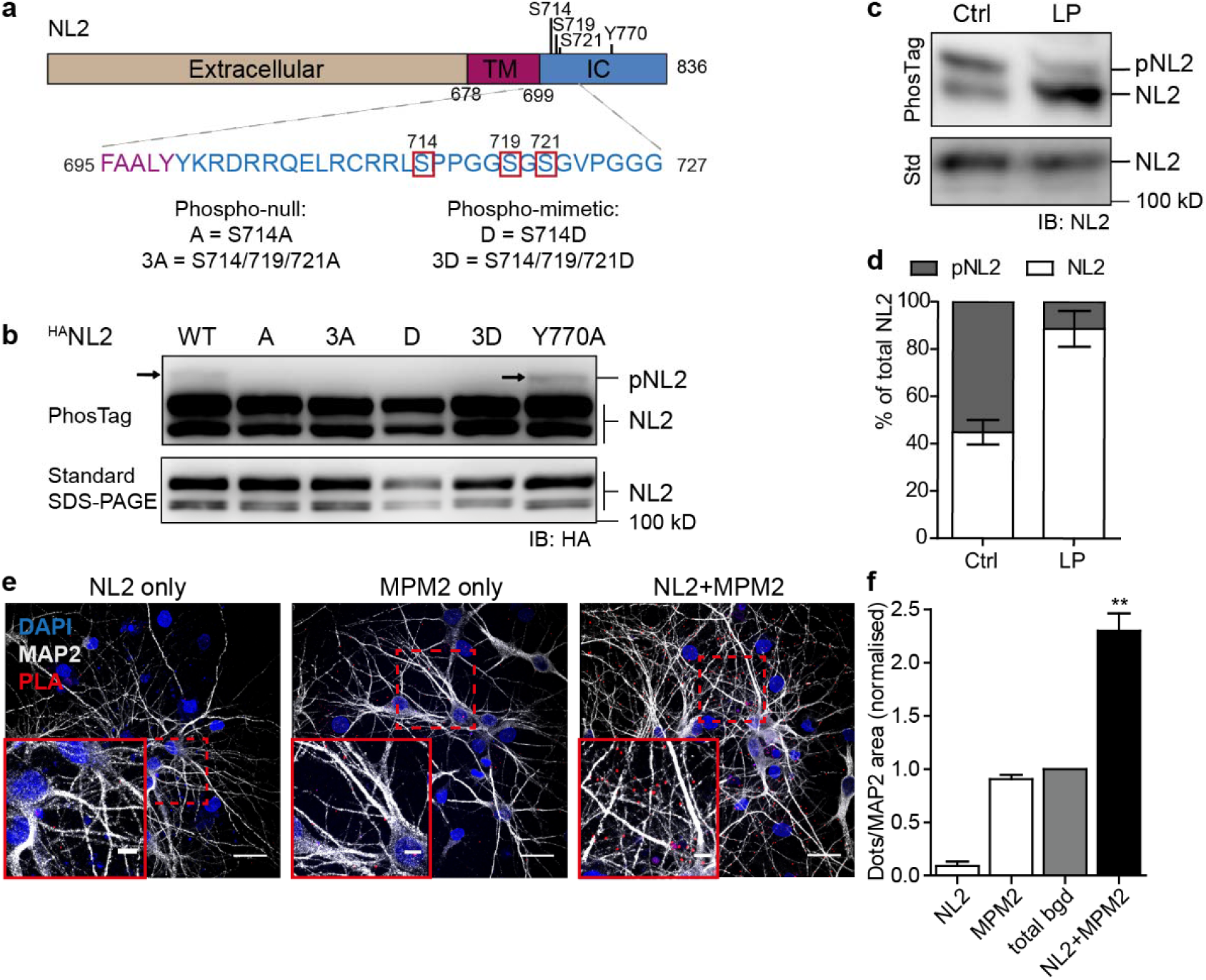
NL2 is constitutively phosphorylated at S714. **a.** Schematic domain structure of murine NL2 showing putative serine phosphorylation sites in the intracellular region. Phospho-null and phospho-mimetic mutations of Serine phosphorylation sites were created as indicated. TM, trans-membrane; IC, intracellular. **b.** Western blot of COS cell lysates expressing WT or mutant ^HA^NL2, separated on standard SDS-PAGE gel (bottom) or SDS-PAGE supplemented with PhosTag (top). Arrows indicate phosphorylated NL2 (pNL2). Molecular weight in kD is indicated on the right. IB, Immunoblot. **c.** Western blot of cortical lysates (DIV8) incubated with buffer (Ctrl) or lambda phosphatase (LP), separated on standard SDS-PAGE gel (Std; bottom) or SDS-PAGE supplemented with PhosTag (top). Molecular weight in kD is indicated on the right. IB, Immunoblot. **d.** Quantification of western blot intensity representing phosphorylated NL2 (pNL2, gray) *vs* unphosphorylated NL2 (NL2, white) as percentage of total NL2; values are mean ± SEM, n=3. **e.** Confocal microscopy images of Proximity Ligation Assay (PLA) in hippocampal cultures using NL2 and MPM2 antibodies and immunostained for MAP2 (gray); PLA signal (red dots) indicates proximity between NL2 and MPM2 antibodies, nuclei are stained with DAPI (blue). Scale bar 25 μm (overview) or 6 μm (zoomed insets). **f.** Quantification of PLA signal between NL2 and MPM2 antibodies, normalised to summed background (total bgd). Values are mean ± SEM; **p<0.01, n=3 experiments (averaged values of 5,14,14 cells per experiment, respectively), two-tailed ratio-paired *t* test between background and NL2+MPM2 (t=11.58).

Given that only a small proportion (5.1 ± 1.2%, n=5) of ^HA^NL2-WT was phosphorylated in COS cells (Fig.1b), we next investigated the extent of phosphorylation of endogenous NL2 in neurons. We incubated cortical lysates with or without lambda phosphatase to dephosphorylate NL2 *in vitro* and analysed the lysates on SDS-PAGE supplemented with PhosTag. Whereas a standard SDS-PAGE reveals one major band for NL2, the PhosTag gel shows two clearly distinguishable bands (Fig.1c). Upon incubation with lambda phosphatase, a strong decrease in intensity of the higher band is observed, whereas intensity of the lower band increases. This suggests that the higher band represents phosphorylated NL2, whereas the lower band represents the non-phosphorylated population, consistent with slower migration of phosphorylated proteins on a PhosTag gel. Quantification of signal intensity reveals that in neurons 55.1 ± 5.2% of NL2 is basally phosphorylated (Fig.1c-d).

To assess whether NL2 phosphorylation at resting state in neurons specifically occurs at S714, we performed a Proximity Ligation Assay (PLA) on rat hippocampal neurons, using an antibody that detects the intracellular domain of NL2, combined with mitotic phosphoprotein monoclonal 2 (MPM2) antibody that specifically recognises phosphorylated S/T-P motifs (Davis et al., 1983) (Fig.1e). Among the serine residues located on the intracellular domain of NL2, S714 is the only serine preceding a proline residue (Fig.1a) and should therefore be specifically recognised by the MPM2 antibody when phosphorylated. PLA signal upon incubation with the individual antibodies was summed to estimate the background. Despite high background for MPM2 alone, we observed a 2.3 ± 0.3-fold increase in signal when both antibodies were combined compared to background (Fig.1f), further supporting our finding that NL2 is phosphorylated at S714 at resting state in neurons.

Taken together, our data suggests that NL2 is constitutively phosphorylated at S714 in neurons.

### PKA mediates rapid phosphorylation of NL2 at S714

To identify factors that can regulate phosphorylation of S714, we investigated a range of treatments that increase neuronal activity by incubating rat neuronal cultures. These factors include 4-aminopyridine (4AP), glutamate, kainic acid and Mg^2+^-free medium, as well as regulators of kinases and phosphatases that commonly play a role in neuronal signalling, such as FK506, forskolin, PMA, and SC79, and analysed the cell lysates on PhosTag gel. Using these experiments, we found that 25 μM forskolin enhances NL2 phosphorylation in cortical and hippocampal cultures (Fig.2a-b; data not shown for other treatments). Quantification showed that 1 hr incubation with forskolin leads to near complete phosphorylation of NL2 (97.7 ± 0.4%). Brief incubation of 5 min leads to a sizeable increase in the amount of phosphorylated NL2 (74.7 ± 5.6%), and 15 min incubation results in 92.2 ± 1.6% of NL2 to be phosphorylated (Fig.2a-b).

**Figure 2:**
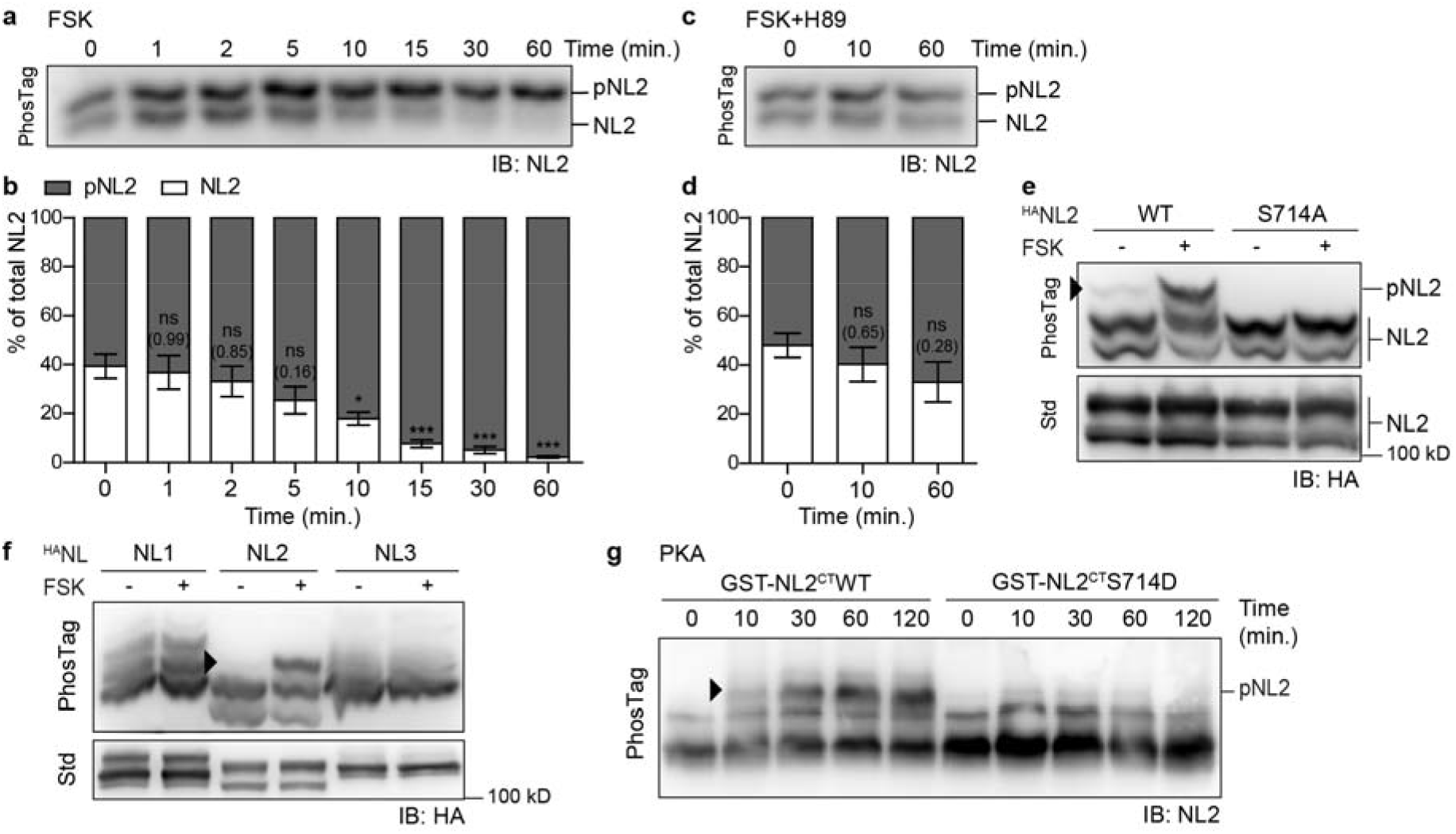
NL2 is phosphorylated at S714 by PKA. **a,c.** Western blots of hippocampal lysates (DIV13) separated on SDS-PAGE supplemented with PhosTag after cell cultures were incubated with 25 μM Forskolin (FSK, **a**), or 25 μM FSK + 50 μM H89 (**c**) for the indicated time points. IB, immunoblot. **b,d**. Quantification of western blot intensity representing phosphorylated NL2 (pNL2, gray) *vs* unphosphorylated NL2 (NL2, white) as percentage of total NL2 upon incubation with FSK (**b**, n=4), or FSK+H89 (**d**, n=3) for the indicated time points. Values are mean ± SEM; ns, not significant (p>0.05, exact p-values are indicated between brackets), *p<0.05, ***p<0.001, 1-way ANOVA with Dunnett’s multiple comparison w.r.t. t=0 (**b:** F(7,24)=11.28, p<0.0001; **d:** F(2,6)=1.21, p=0.36). **e,f.** Western blots of COS cell lysates overexpressing WT or S714A ^HA^NL2 (**e**) or ^HA^NL1, ^HA^NL2 or ^HA^NL3 (**f**) separated on standard SDS-PAGE gel (Std; bottom) or SDS-PAGE supplemented with PhosTag (top) after cultures were treated with FSK (+) or DMSO (-). Arrowheads indicate phosphorylated ^HA^NL2. Molecular weight in kD is indicated on the right. IB, Immunoblot. **g**. Western blot of purified GST-NL2^CT^WT or GST-NL2^CT^S714D incubated with purified catalytic subunit of PKA (50 U/μl) for the indicated time points, separated on SDS-PAGE supplemented with PhosTag. The arrowhead indicates phosphorylated GST-NL2^CT^WT upon incubation with PKA.

Forskolin activates cAMP-dependent protein kinase (PKA) through increasing intracellular levels of cAMP (Nguyen and Woo, 2003). Indeed, the site S714 in NL2 is compatible with the consensus sequence for PKA-induced phosphorylation, RRXS/TY (Fig.1a). To verify whether forskolin-induced phosphorylation of NL2 is mediated by PKA, we incubated hippocampal cultures simultaneously with forskolin and the PKA-specific inhibitor H89. Quantification of the lysates on PhosTag gel (Fig.2c-d) showed no significant change in the amount of phosphorylated NL2, even after 1 hr of incubation, confirming that NL2 phosphorylation is dependent on PKA.

To verify whether forskolin-induced phosphorylation of NL2 occurs at S714, we incubated COS cells expressing WT or S714A ^HA^NL2 with forskolin. Analysis of cell lysates on PhosTag gel revealed that incubation with forskolin increased levels of phosphorylated ^HA^NL2-WT, whereas no change was seen for ^HA^NL2-S714A (Fig.2e). This confirms that forskolin enhances phosphorylation of NL2 specifically at S714. Interestingly, despite the PKA-consensus sequence being conserved in other Neuroligin isoforms, we find that forskolin-induced phosphorylation is unique for NL2: when incubating COS cells overexpressing ^HA^NL1, ^HA^NL2 or ^HA^NL3 with forskolin, we observe a change in banding pattern on PhosTag gel only for ^HA^NL2-containing lysates (Fig.2f).

Finally, to verify *in vitro* whether phosphorylation of NL2 is mediated by PKA, we fused the intracellular C-terminal tail of NL2 to GST (GST-NL2^CT^) and incubated purified GST-NL2^CT^WT or phospho-mutant GST-NL2^CT^S714D with purified PKA (50 U/μl; Fig 2g). Analysis on PhosTag gel shows an increase in phosphorylation over time for WT but not S714D GST-NL2^CT^, further confirming that PKA phosphorylates NL2 at S714.

Taken together, these results show that PKA mediates rapid phosphorylation of NL2 at S714, and that this pathway can be induced by forskolin.

### Phosphorylation of NL2 leads to its dispersion from the synapse

Given the role of phosphorylation in regulating protein interactions and trafficking, we investigated whether forskolin-mediated NL2 phosphorylation affected its synaptic stabilisation and surface distribution. First, we investigated how phosphorylation affects real-time membrane diffusion of NL2 using single molecule tracking with quantum dots (QDs). Hippocampal neurons were co-transfected with ^HA^NL2-WT or ^HA^NL2-S714A and Gephyrin-FingR, a GFP-conjugated intracellularly expressed IgG-like domain that specifically recognises gephyrin clusters (Gross et al., 2013). This allows for simultaneous identification of inhibitory synapses via Gephyrin-FingR, and QD tracking of surface-exposed ^HA^NL2 molecules using anti-HA primary and QD-conjugated secondary antibodies (Fig.3a). WT and S714A ^HA^NL2 diffuse dynamically on the plasma membrane and localise to synaptic and extrasynaptic microdomains (Fig.3b). In addition, some ^HA^NL2s exchanged localisation between synaptic and extrasynaptic areas, suggesting that ^HA^NL2s are recruited to inhibitory synapses by lateral diffusion. As expected, from the mean square displacement (MSD) plots, synaptic ^HA^NL2s diffuse slower and were more confined compared to extrasynaptic ^HA^NL2s (Fig.3c-f; Median synaptic diffusion coefficient, *D*_*synaptic*_ 0.040 μm^2^s^−1^, *D*_*extrasynaptic*_ 0.080 μm^2^s^−1^; Median confinement area *CA*, *CA*_*synaptic*_ 0.29 μm^2^; *CA*_*extrasynaptic*_ 0.33 μm^2^). These parameters suggest that ^HA^NL2 is showing similar dynamics to that expected for endogenous NL2.

**Figure 3:**
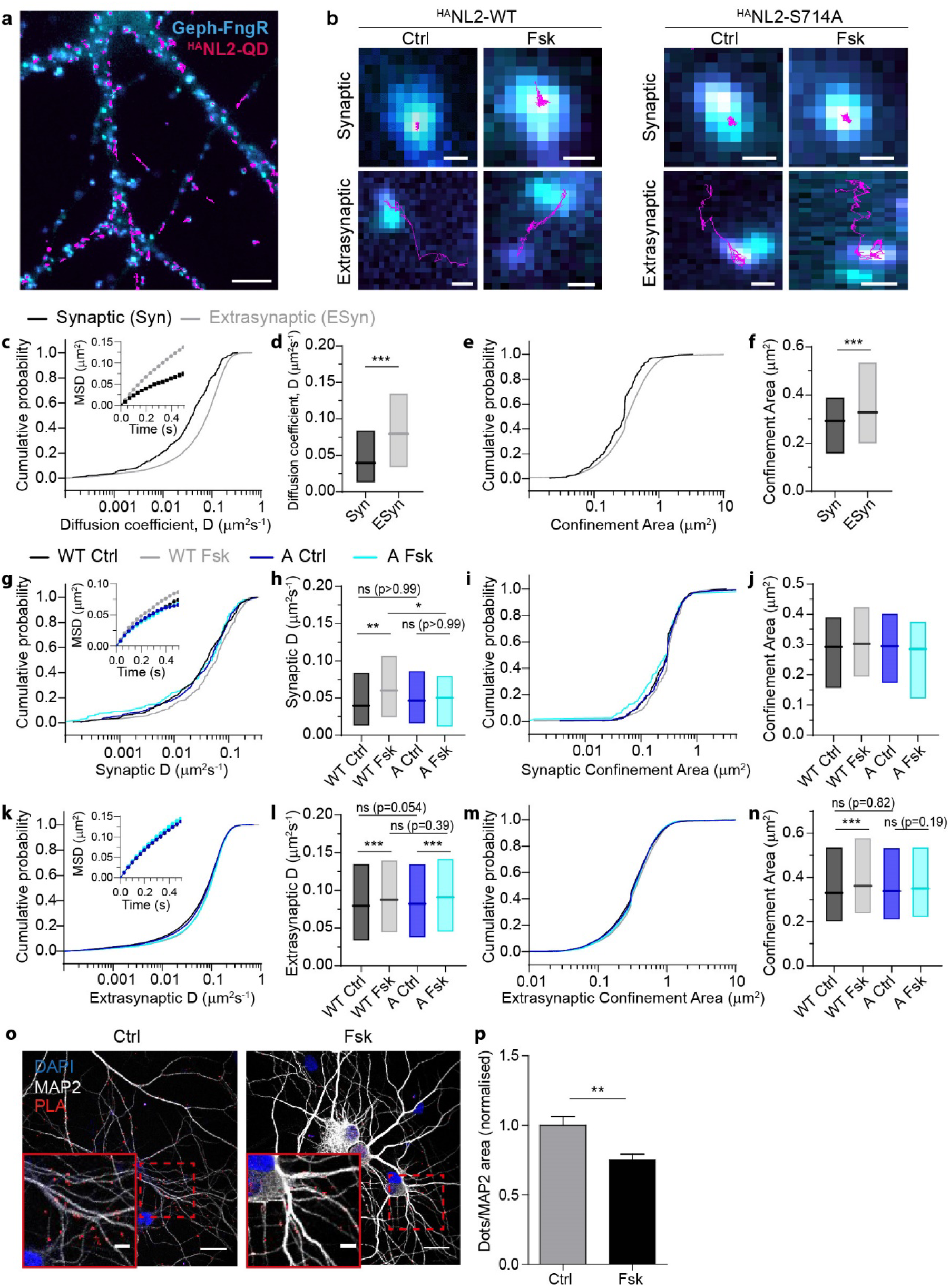
Phosphorylation of NL2 at S714A causes its synaptic dispersion. **a.** Fluorescence microscopy image of live hippocampal culture (DIV13), co-expressing gephyrin-FingR (Geph-FngR, cyan) and ^HA^NL2 (magenta, QD trajectories summed from 300 frames at 33 Hz). Scale bar 5 μm. **b**. Example trajectories of overexpressed WT (left panel) or S714A (right panel) ^HA^NL2 (magenta) treated with DMSO (Ctrl) or 25 μM Forskolin (Fsk); trajectories were classified as synaptic (top) or extrasynaptic (bottom) based on colocalization with Geph-FngR (cyan). Scale bar 0.5 μm. **c,e.** Cumulative probability plot for diffusion coefficients, D (**c**, main graph) or Confinement Area (**e**), and plot of mean square displacement (MSD) showing increased confinement of synaptic receptors (**c**, inset) for synaptic (black, n=277) and extrasynaptic (gray, n=13962) trajectories of ^HA^NL2-WT. **d,f.** Box plot of the median and inter-quartile range (IQR, 25%– 75%) of D values (**d**) or Confinement Area (**f**) for synaptic (Syn) and extrasynaptic (Esyn) trajectories of ^HA^NL2-WT; ***p<0.001, two-tailed Mann-Whitney test. **g,i.** Cumulative probability plot for diffusion coefficients (D) (**g**, main graph) or Confinement Area (**i**), and plot of mean square displacement (MSD) (**g**, inset) for synaptic trajectories of ^HA^NL2-WT and ^HA^NL2-S714A, upon treatment with DMSO (Ctrl) or Forskolin (Fsk); WT Ctrl, n=277; WT Fsk n=297; A Ctrl, n=343; A Fsk, n=174. **h,j.** Box plot of the median and IQR of D values (**h**) or Confinement Area (**j**) for synaptic trajectories of ^HA^NL2-WT and ^HA^NL2-S714A, upon treatment with DMSO (Ctrl) or Forskolin (Fsk); ns, not significant (p>0.05, exact p-values are indicated between brackets), *p<0.05, **p<0.01, ***p<0.001; Kruskal-Wallis tests with Dunn’s multiple comparison (**h:** overall p=0.0024; **j:** overall p=0.068). **k-n**. As for g-j but displaying extrasynaptic trajectories; WT Ctrl, n=13962; WT Fsk n=10616; A Ctrl, n=13317; A Fsk, n=10699 (**l:** overall p<0.0001; **n:** overall p<0.0001). **o.** Confocal microscopy images of Proximity Ligation Assay (PLA) in hippocampal cultures using NL2 and Gephyrin antibodies and immunostained for MAP2 (gray); PLA signal (red dots) indicates proximity between NL2 and Gephyrin antibodies upon treatment with DMSO (Ctrl) or Forskolin (Fsk); nuclei are stained with DAPI (blue). Scale bar 25 μm (overview) or 6 μm (zoomed insets). **p.** Quantification of normalised PLA signal between NL2 and Gephyrin antibodies. Values are mean ± SEM; Ctrl n=41, Fsk n=48 cells from 4 independent cultures; **p<0.01, two-tailed unpaired *t* test (t=3.37).

The effect of forskolin-mediated PKA phosphorylation of S714 on mobility of NL2 was studied by incubating neurons in vehicle (DMSO, Ctrl) or 25 μM forskolin (Fsk) for 5 min before imaging. Synaptic trajectories of ^HA^NL2-WT show a clear increase of *D* upon treatment with forskolin (*D*_*Ctrl*_ 0.040 μm^2^s^−1^, *D*_*Fsk*_ 0.060 μm^2^s^−1^), whereas ^HA^NL2-S714A trajectories are unaffected (*D*_*Ctrl*_ 0.047 μm^2^s^−1^, *D*_*Fsk*_ 0.050 μm^2^s^−1^) (Fig.3g-h). Confinement area remained unaffected (Fig.3i-j; WT: *CA*_*Ctrl*_ 0.29 μm^2^, *CA*_*Fsk*_ 0.30 μm^2^; S714A: *CA*_*Ctrl*_ 0.29 μm^2^, *CA*_*Fsk*_ 0.29 μm^2^), although individual comparison showed a significant increase in confinement area for ^HA^NL2-WT upon addition of forskolin (p=0.044, two-tailed Mann-Whitney test). By contrast, forskolin only marginally increased extrasynaptic lateral diffusion of ^HA^NL2-WT (*D*_*Ctrl*_ 0.080 μm^2^s^−1^, *D*_*Fsk*_ 0.088 μm^2^s^−1^; *CA*_*Ctrl*_ 0.33 μm^2^, *CA*_*forskolin*_ 0.36 μm^2^) and ^HA^NL2-S714A (*D*_*Ctrl*_ 0.082 μm^2^s^−1^, *D*_*Fsk*_ 0.091 μm^2^s^−1^; *CA*_*Ctrl*_ 0.34 μm^2^, *CA*_*forskolin*_ 0.35 μm^2^s) (Fig.3k-n), suggesting that WT and S714A ^HA^NL2 are minimally dispersed extrasynaptically. Thus, phosphorylation does not play a major role in extrasynaptic lateral diffusion of NL2. Indeed, *D* of forskolin-treated WT and S714A ^HA^NL2 extrasynaptic receptors were not significantly different while at the synapse WT ^HA^NL2 mobility was higher compared to S714A due to phosphorylation. Together these results suggest that upon phosphorylation at S714, synaptic NL2 are dispersed from synapses.

To investigate whether phosphorylation-induced dispersal of NL2 from the synapse coincides with a reduced proximity to gephyrin, we investigated the interaction between endogenous NL2 and gephyrin using PLA (Fig.3o-p). Using this method, we observed a 25 ± 0.4% decrease in PLA signal upon forskolin treatment, representing decreased proximity between NL2 and gephyrin, presumably by decreased interaction.

Thus, our results are in accord with a destabilisation of NL2 from the inhibitory postsynaptic scaffold upon phosphorylation.

### Phosphorylation of NL2 reduces surface expression and affects inhibitory signalling

Dispersal of NL2 from the synapse and reduced interaction of NL2 with gephyrin upon PKA-induced phosphorylation suggests that phosphorylation may affect surface distribution of NL2. We used surface protein biotinylation to investigate the effect of PKA-mediated NL2 phosphorylation on cell surface levels and found that phosphorylation resulted in a 40.4 ± 7.7% decrease in surface-exposed NL2 (Fig.4a-b). This suggests that phosphorylation of NL2 at S714 may enhance its internalisation. To further investigate the role of residue S714 in internalisation, we overexpressed ^HA^NL2-WT, ^HA^NL2-S714A or ^HA^NL2-S714D in hippocampal neurons and assessed basal internalisation using antibody feeding, making use of the extracellular HA-tag to live-label NL2 with antibody and allow for it to internalise (Fig.4c). Quantification of internalised *vs* surface ^HA^NL2 reveals that internalisation of both ^HA^NL2-S714A and ^HA^NL2-S714D is reduced compared to ^HA^NL2-WT, with S714D being less impaired than S714A (Fig.4d). Thus, an intact phosphorylation site at S714 is required for internalisation of NL2, suggesting internalisation is phosphorylation dependent. Furthermore, this suggests that the D-mutant functions as a partial phospho-null rather than full phospho-mimetic mutant.

**Figure 4:**
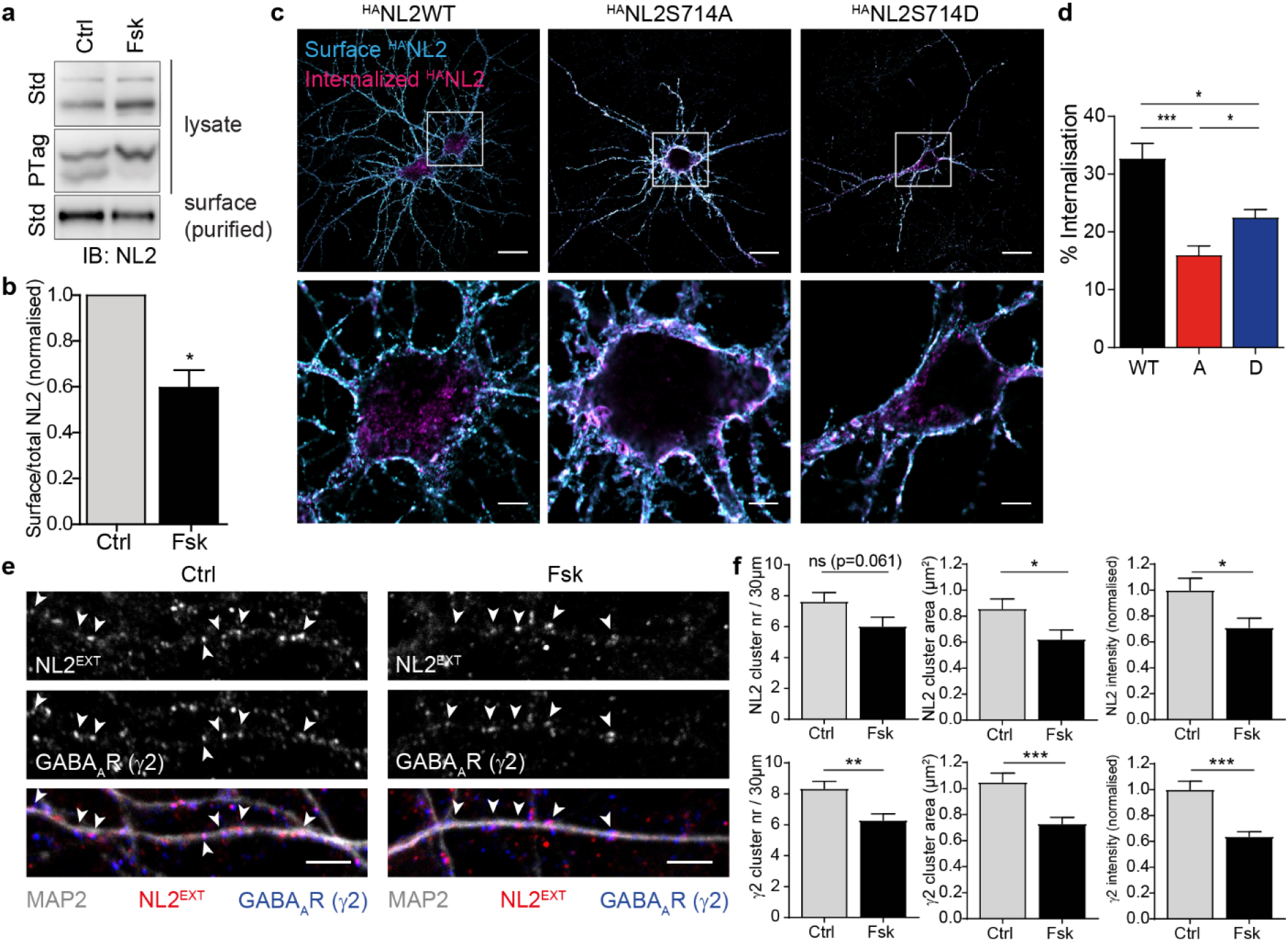
Phosphorylation of NL2 at S714A reduces NL2 and GABA_A_R surface levels. **a.** Western blot of cortical lysates (DIV8; top, middle) and purified surface protein (bottom) upon treatment with DMSO (Ctrl) or Forskolin (Fsk), separated on standard SDS-PAGE gel (Std; top, bottom) or SDS-PAGE supplemented with PhosTag (PTag, middle). **b.** Quantification of normalised western blot signal intensity of surface NL2 as a percentage of total NL2; n=3, *p<0.05, paired two-tailed *t* test (t=5.25). **c**. Confocal microscopy images of antibody feeding in hippocampal neurons (DIV13) overexpressing ^HA^NL2WT (WT, left), ^HA^NL2S714A (A, middle) or ^HA^NL2S714D (D, right), showing surface-exposed (cyan) and internalised ^HA^NL2 (magenta). Scale bar 25 μm (overview, top) or 5 μm (zoom, bottom). **d**. Quantification of internalisation for WT (n=33), S714A (A, n=28) and S714D (D, n=44) ^HA^NL2 in hippocampal neurons. Values are mean ± SEM, *p<0.05, ***p<0.001, Kruskal-Wallis test with Dunn’s multiple comparison test (overall p<0.0001). **e.** Confocal microscopy images of 30 μm dendritic sections of hippocampal neurons (DIV13), immunostained for surface NL2 (NL2^EXT^, red), GABA_A_R subunit γ2 (blue) and MAP2 (gray), and treated with DMSO (Ctrl, left panel) or Forskolin (Fsk, right panel). Arrowheads indicate synaptic clusters of co-localising NL2^EXT^ and γ2. Scale bar 4 μm. **f.** Quantification of cluster number (left), cluster area (middle) and normalised cluster staining intensity (right) of surface NL2 (top) and γ2 (bottom), treated with DMSO (Ctrl, n=19) or Forskolin (Fsk, n=20); ns, not significant (p>0.05, exact p-values are indicated between brackets), *p<0.05, **p<0.01, ***p<0.001, unpaired two-tailed *t* tests.

To investigate how internalisation of NL2 upon phosphorylation affects the concentration of synaptic GABA_A_Rs, we quantified surface staining of synaptic clusters in hippocampal neurons treated with DMSO or forskolin, using antibodies recognising extracellular regions of the GABA_A_R γ2 subunit and endogenous NL2, respectively (Fig.4e-f). Although the reduction in surface NL2 cluster number did not reach significance, we found a significant reduction in NL2 and γ2 cluster area and fluorescence intensity, suggesting a decrease in concentration of both markers from synapses. This implies that loss of synaptic NL2 coincides with destabilisation of GABA_A_Rs at the inhibitory synapse, consistent with data from our recent study (Halff et al., 2019).

Finally, to test whether phosphorylation of NL2 at S714, and altered levels of synaptic NL2 and GABA_A_Rs, impacts on inhibitory signalling, we performed electrophysiology, measuring sIPSCs from cultured hippocampal neurons overexpressing either WT, S714A, or S714D ^HA^NL2 (Fig.5a). We found no change in sIPSC frequency with either mutant compared to ^HA^NL2WT, suggesting presynaptic release remained unaffected (Fig.5b). In contrast, both S714A and S714D overexpression increased the sIPSCs amplitude compared to WT ^HA^NL2 (Fig.5a,c-d), with S714D showing a smaller increase than S714A. This suggests that the inability of NL2 to be phosphorylated, and consequently its increased stability at the synapse and reduced internalisation, results in enhanced inhibitory signalling, possibly indirectly through increased stabilisation of postsynaptic complexes incorporating GABA_A_Rs. Furthermore, we find that S714D has an intermediate phenotype, as was also observed in the internalisation essay (Fig.4d), thus serving as a partial phospho-null rather than phospho-mimetic mutation. Interestingly, analysis of sIPSC amplitude histograms revealed three gaussian-distributed peaks for WT and S714D whereas a fourth peak was observed for S714A at −1771 pA (Fig.5e). Moreover, the high-amplitude peaks were notably increased for NL2 mutants compared to WT, with the shift being more pronounced for S714A than S714D. This suggests that the overall increase in sIPSC amplitude (Fig.5d) was mainly caused by the occurrence of larger GABAergic synapses upon overexpression of the S714A mutant. Overall, these data suggest that interfering with NL2 phosphorylation leads to enhanced inhibitory signalling, which may be through stabilisation of postsynaptic components at the inhibitory synapse.

**Figure 5:**
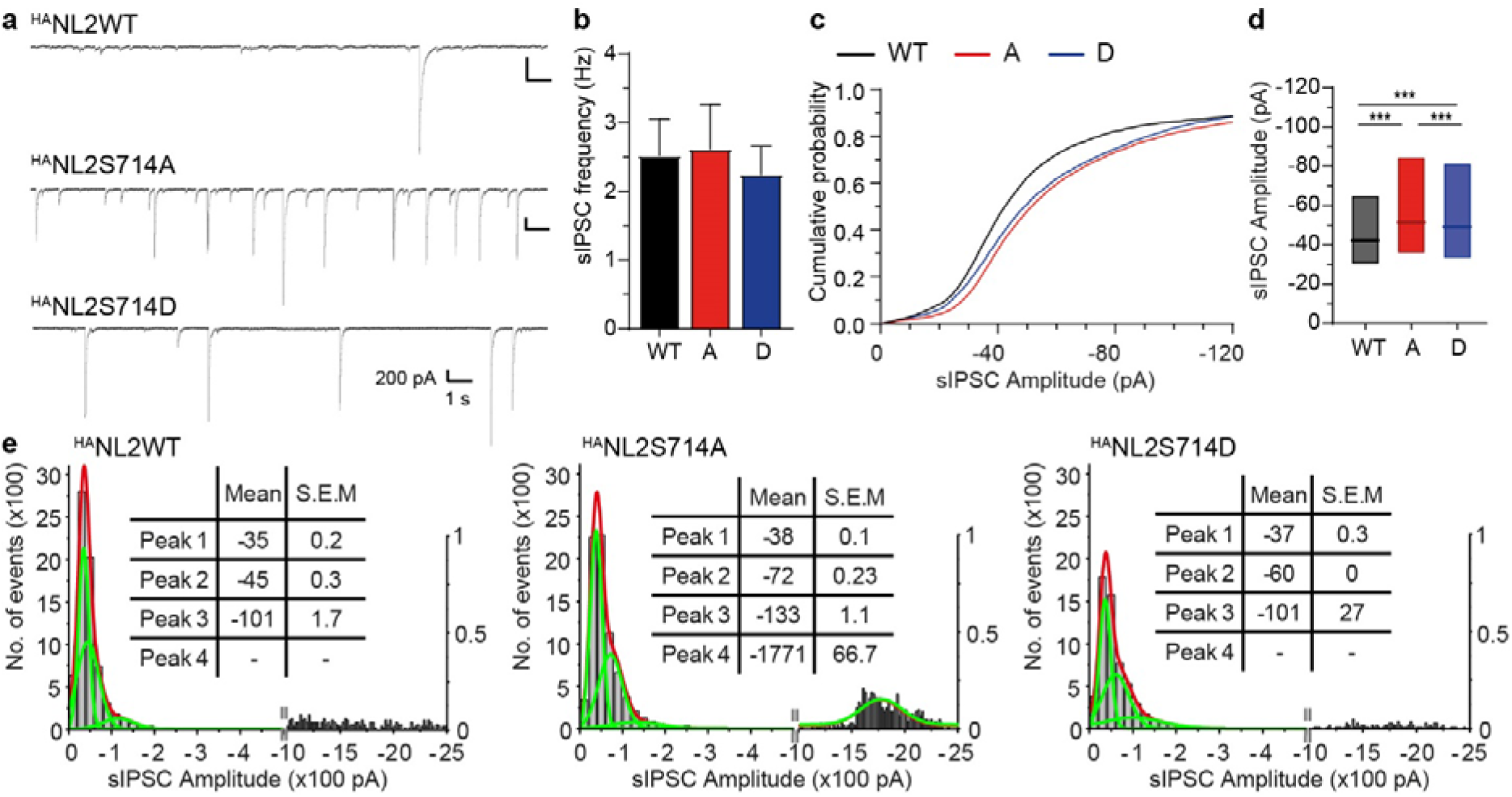
Phosphorylation of NL2 at S714A affects inhibitory signalling. **a.** Representative sIPSC patch-clamp recordings from hippocampal cultures expressing ^HA^NL2WT, ^HA^NL2S714A or ^HA^NL2S714D combined with eGFP. **b.** Pooled data of sIPSC frequencies; WT, n=16; A, n=18; D, n=15; one-way Anova (F(2,46)=0.11, overall p=0.89). **c,d.** Cumulative probability plot (**c**) and box plot of median and Inter-quartile range (25-75%, **d**) of sIPSC amplitudes; ***p<0.001, Kruskal-Wallis test with Dunn’s multiple comparison (overall p<0.0001). **e.** Histograms of sIPSC amplitudes showing individual Gaussian-distributed peaks at distinct amplitudes (green lines and table) compared to the overall distribution (red line) upon expression of ^HA^NL2 WT and mutants.

Taken together, these experiments suggest that NL2 surface levels are modulated by PKA-mediated phosphorylation at S714, consequently altering synaptic levels of GABA_A_Rs and impacting on inhibitory signalling strength.

## Discussion

Neuroligin-2 (NL2) is essential for the maintenance and stabilisation of inhibitory synapses, both through recruitment of a gephyrin scaffold that stabilises GABA_A_Rs at the synapse, and through trans-synaptic interaction with Neurexins to ensure correct positioning of the GABAergic postsynaptic domain (Craig and Kang, 2007, Sudhof, 2008, Bemben et al., 2015). Modulation of synaptic NL2 levels contributes to the regulation of inhibitory signalling (Binda et al., 2019, Halff et al., 2019), however, the mechanisms regulating surface levels and synaptic stabilisation of NL2 remained poorly understood. In the current study, we identify a role for cAMP-dependent protein kinase (PKA) in regulating these processes. We find that PKA directly phosphorylates NL2 at S714 and provide evidence that PKA-mediated phosphorylation at this site causes its rapid dispersal from inhibitory synaptic clusters. Enhanced phosphorylation leads to reduced surface levels of NL2 and, consequently, reduces the concentration of synaptic GABA_A_Rs. Conversely, abolishing NL2 phosphorylation by mutation of S714 reduces its internalisation and enhances inhibitory signalling. Thus, PKA-mediated phosphorylation of NL2 contributes to the regulation of synaptic inhibition by modulating NL2 synaptic stability and surface levels (Fig.6a).

**Figure 6:**
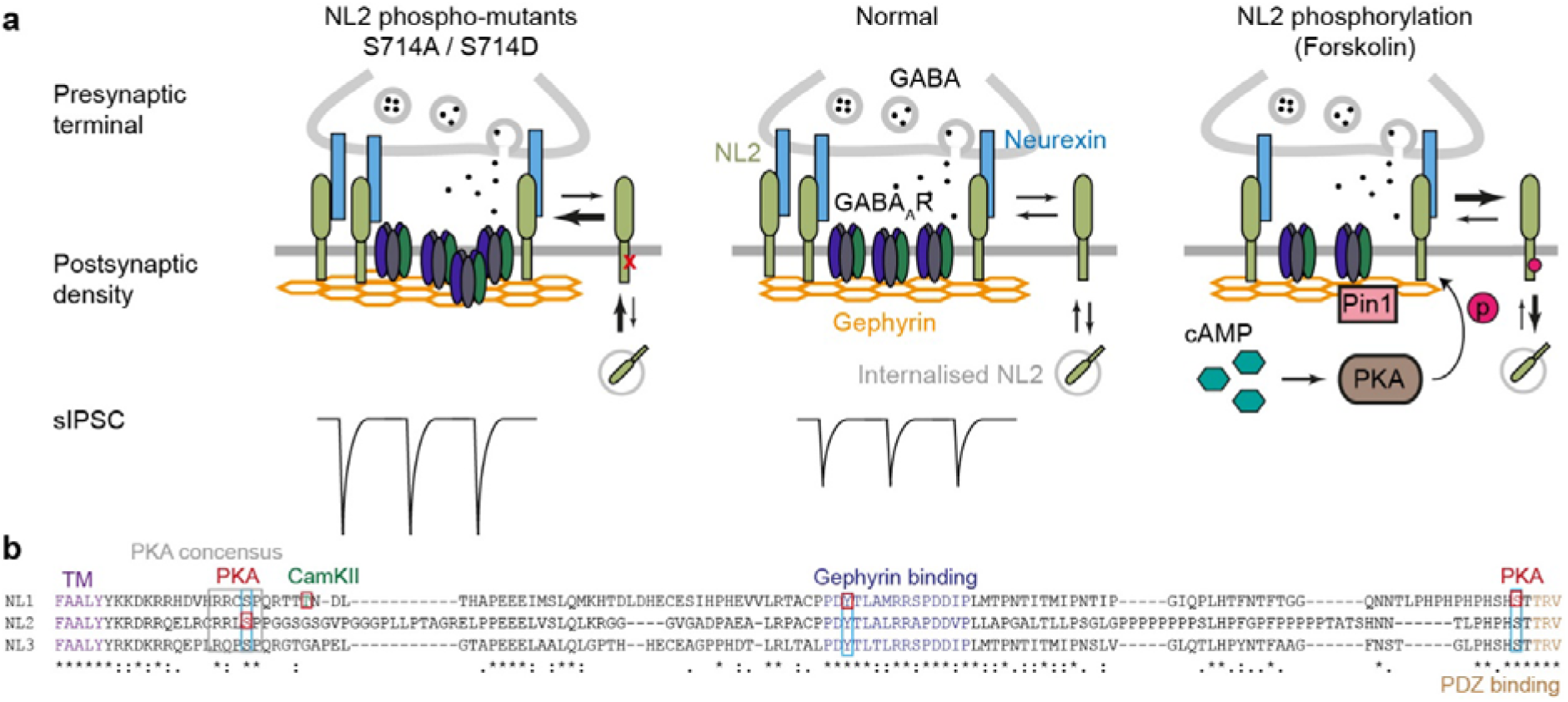
Unique PKA-mediated phosphorylation site in NL2 mediates its surface and synaptic stability. **a.** Schematic representation of the proposed molecular model showing the effects of NL2 phosphorylation at S714. Forskolin (right-hand side) activates PKA through increasing intracellular cAMP levels; as a result, NL2 is phosphorylated, recruits Pin1, disperses from the synapse and is internalised. Either through loss of synaptic NL2 directly or through Pin1 activity, other components of the inhibitory postsynaptic site are destabilised, including loss of synaptic GABA_A_Rs. Abolishing NL2 phosphorylation through mutation of S714 (left-hand side) reduces internalisation and stabilises NL2 at the synapse, leading to increased GABA_A_R concentration and enhanced sIPSC amplitude. **b.** Sequence alignment of mouse NL1, NL2 and NL3 showing the position of the NL2-S714 PKA phosphorylation site and conserved consensus sequence (gray box), as well as characterised phosphorylation sites in other NL isoforms (red squares). Light blue boxes indicate conservation of these phosphorylation sites. TM, trans-membrane.

Phosphorylation of NL2 and its subsequent dispersion from the synapse may be an important mechanism in rapidly responding to neuronal activity, fine tuning inhibitory signalling, and maintaining the E/I balance. Lateral diffusion is a major mechanism for rapid (in seconds) recruitment of receptors and trans-membrane proteins into synapses as well as for their dispersal from the synapse (Bannai et al., 2009, Choquet and Triller, 2013). Our data from single molecule tracking and electrophysiology combined suggests that NL2 phosphorylation contributes to rapid downregulation of inhibitory signalling as a first step in destabilising the GABAergic synapse and, conversely, stabilisation of unphosphorylated NL2 at the synapse may lead to an increase in synapse size (Fig.6a).

Analysis of cortical lysates revealed that 50-60% of endogenous NL2 is phosphorylated in neuronal cultures (Fig.1c-d). Whether this population mostly resides intracellularly or remains at the cells surface extrasynaptically remains to be investigated. Surface biotinylation upon PKA activation showed only a partial reduction of NL2 from the cell surface (Fig.4a-b), despite near-complete phosphorylation at this time point (Fig.2a-b). This suggests the majority of phosphorylated NL2 remains at the cell surface, where it can be dephosphorylated and rapidly re-join the synapse. Conversely, it could also be that, whereas phosphorylation induces NL2 to be internalised, it gets replaced rapidly by membrane insertion of non-phosphorylated NL2. Long-term incubation (1 hr) with forskolin, as used in our biotinylation experiment, may push the balance towards enhanced internalisation. In either scenario, NL2 phosphorylation provides a putative mechanism for brief downregulation of inhibitory synapses, which can be quickly restored upon dephosphorylation of extrasynaptic NL2, or membrane reinsertion followed by rapid lateral diffusion into the synapse.

Given that mutations in Neuroligins associated with neurological and cognitive disorders have been suggested to affect their surface levels and trafficking (Sudhof, 2008, Bemben et al., 2015, Chih et al., 2004), a better understanding of the regulation of Neuroligin trafficking may form an important basis towards improved insight into the molecular basis of these disorders. NL2 can be internalised to endosomes (Kang et al., 2014), and we recently showed that sorting-nexin 27 (SNX27) regulates recycling of internalised NL2, thereby modulating NL2 surface levels (Halff et al., 2019). Our current study demonstrates that phosphorylation of NL2 at S714 controls synaptic stability of NL2 and plays a role in its internalisation. The loss of NL2 from the synapse, either by increased phosphorylation or blocking recycling, reduces synaptic GABA_A_Rs, and stabilising NL2 at the synapse by abolishing phosphorylation or increasing recycling enhances inhibitory signalling.

Our data are consistent with and complimentary to an earlier study, which showed that NL2 requires an intact S714 phosphorylation site to recruit the peptidyl-prolyl isomerase Pin1, and that Pin1 activity negatively regulates interaction between NL2 and gephyrin (Antonelli et al., 2014). In Pin1^−/−^ animals, where downstream effects of NL2 phosphorylation are abolished, the number of GABA_A_Rs at inhibitory synapses was found to be enhanced, leading to an increased amplitude of sIPSCs. Consistently, we find that overexpression of ^HA^NL2-S714A and ^HA^NL2-S714D, which cannot be phosphorylated, enhances inhibitory signalling. This is also in agreement with another recent study, which found that overexpression of either ^HA^NL2-S714A or ^HA^NL2-S714D in a Neuroligin1-3-null background in biolistically transfected brain slices restored inhibitory signalling (Nguyen et al., 2016). We furthermore demonstrate the reverse of this mechanism, where increasing NL2 phosphorylation levels lead to its dispersal from the synapse and reduces synaptic GABA_A_R clusters. All these results combined show that NL2 phosphorylation, either directly through dispersal from the synapse, or indirectly through Pin1 activation and other downstream effects such as conformational changes and altered interactions between inhibitory postsynaptic proteins, impacts on signalling at the inhibitory synapse (Fig.6a).

We noted that PKA-mediated phosphorylation at S714 is unique for NL2, despite the consensus sequence being conserved in other Neuroligin isoforms (Fig.2f, 6b). Downstream of S714, NL2 displays a unique motif, which may contain elements that promote PKA binding or, conversely, a different conformation in other NLs may hinder PKA-mediated phosphorylation at this site. Just downstream of the PKA site in NL2, NL1 was shown to be uniquely phosphorylated at T739 by CamKII in response to synaptic activity (Bemben et al., 2014), enhancing its surface expression and strengthening postsynaptic excitatory signalling. A conserved Tyrosine phosphorylation site in NL1 (Y782) and NL2 (Y770) regulates their preferential specialisation for excitatory synapses and PSD95, or inhibitory synapses and gephyrin, respectively (Giannone et al., 2013, Poulopoulos et al., 2009, Letellier et al., 2020). PKA-mediated phosphorylation of NL1 at S839, a residue directly preceding the highly conserved C-terminal PDZ-binding motif, was shown to abolish its interaction with PSD95 and enhance NL1 internalisation (Jeong et al., 2019). A similar mechanism involving this residue has not yet been identified for other NL isoforms. Thus, despite relatively high sequence conservation of the Neuroligin intracellular domains, there is accumulating evidence that differential phosphorylation of Neuroligin isoforms contributes to their specialisation at specific synapses and isoform-specific activity regulation (Jeong et al., 2017).

What neuromodulatory signalling pathways result in PKA-mediated phosphorylation of NL2, as well as which phosphatases and pathways mediate dephosphorylation and, hypothetically, the stabilisation of synaptic NL2, remains to be established. Many neuronal processes, including signalling by a wide range of hormones and neurotransmitters, increase intracellular cAMP levels, subsequently activating PKA. PKA has been shown to be important for synaptic plasticity (Nguyen and Woo, 2003). GABA_A_R subunits β1 and β3 contain PKA phosphorylation sites, where phosphorylation of the β1 subunit inhibits GABAergic signalling, whereas phosphorylation of β3 has the opposite effect (Nakamura et al., 2015, McDonald et al., 1998). The scaffold protein gephyrin is also heavily regulated by phosphorylation, containing at least 26 putative phospho-sites, which regulate its ability to oligomerise and stabilise GABA_A_Rs (Zacchi et al., 2014, Tyagarajan and Fritschy, 2014). Specifically, one of these sites has been shown to be phosphorylated by PKA (Flores et al., 2015, Battaglia et al., 2018), which was suggested to affect GABA_A_R confinement at the synapse. The role of PKA-mediated phosphorylation of NL2 was not investigated in these studies. Excitatory targets of PKA include subunits of the AMPA and NMDA receptors (Wang et al., 2014). Thus, the net effect of PKA activation on neuronal activity may be a summation of many factors including compartmentalisation of PKA and accessibility of its binding sites. Due to the complex interplay between these pathways involving PKA we cannot predict how the effect of PKA-activation on other synaptic components plays a role in our experiments, or how NL2 phosphorylation and destabilisation contributed to effects of PKA activation on inhibitory signalling reported in earlier studies. However, all data taken together, we have shown a clear and direct connection between PKA-mediated phosphorylation of NL2, its molecular surface dynamics and surface levels, and inhibitory signalling. This research thus furthers our understanding of how the function of NL2 is regulated, and how this modulates inhibitory signalling strength.

## Acknowledgements

We would like to thank Dr. David F. Sheehan for initial guidance on single particle tracking. This work was supported by a grant to JTK from the BBSRC (BB/I00274X/1). EFH received funding from the European Union’s Horizon 2020 research and innovation programme under the Marie Skłodowska-Curie grant agreement No.661733. SH and TGS are supported by an MRC programme grant and Wellcome Trust collaborative grant. SH also has an International Rett Syndrome Foundation Fellowship (3606).

## Author contributions

Conceptualisation & Supervision: JTK; Methodology: EFH, JTK, SH; Formal analysis: EFH, SH, TGS; Investigation and Visualisation: EFH, SH; Resources: JTK, TGS; Funding Acquisition: JTK, EFH; Writing – Original Draft: EFH, SH; Writing – Review & Editing: all authors

## Materials & Correspondence

Correspondence and material requests should be addressed to

## Experimental Procedures

### Animals

All procedures for the care and treatment of animals were in accordance with the Animals (Scientific Procedures) Act 1986, and had full Home Office ethical approval. Animals were maintained under controlled conditions (temperature 20 ± 2°C; 12h light-dark cycle), were group housed in conventional cages and had not been subject to previous procedures. Food and water were provided ad libitum. Wild-type E18 Sprague-Dawley rats were generated as a result of wild-type breeding; pregnant females were single housed. Embryos of either sex were used for generating primary neuronal cultures.

### Primary Hippocampal Culture and transfection

Cultures of hippocampal and cortical neurons were prepared from E18 WT Sprague-Dawley rat embryos of either sex as previously described (Davenport et al., 2019, Hannan et al., 2020). In brief, rat hippocampi and cortices were dissected from embryonic brains in ice-cold HBSS (GIBCO) supplemented with 10mM HEPES. Dissected hippocampi were incubated in the presence of 0.25% trypsin and 5 Units/ml DNase for 15 mins at 37°C, washed twice in HBSS with HEPES, and triturated to a single cell suspension in attachment media (MEM (GIBCO) containing 10% horse serum, 10 mM sodium pyruvate, and 0.6% glucose) using a fire-polished glass Pasteur pipette. Dissociated cortical neurons were plated in attachment media on poly-L-lysine (PLL) coated dishes at a density of 2×10^6^ cell/6 cm dish. Dissociated hippocampal neurons were plated in attachment media on PLL coated coverslips at a density of 5×10^5^ cells/6 cm dish (PLA, cluster analysis, antibody feeding), 1.5×10^6^ cells/6 well dish (single particle tracking, electrophysiology), or PLL coated dishes at a density of 2×10^6^ cells/12 well dish (forskolin timecourse treatments). After 6 hr, serum-containing medium was replaced with Neurobasal medium (GIBCO) containing 2% B-27 (GIBCO), 2 mM glutaMAX (GIBCO), 100 U/ml penicillin and 100 μg/ml streptomycin. Cultures were maintained at 37°C in humidified atmosphere with 5% CO_2_. Hippocampal neurons were transfected using a calcium phosphate method (Hannan et al., 2020) or Lipofectamine 2000 (Invitrogen) at DIV7 for cluster analysis and electrophysiology experiments, or at DIV10 for antibody feeding and single particle tracking, and maintained until DIV13-15.

### COS-7 cell culture and transfection

COS-7 cells were maintained in DMEM (GIBCO), supplemented with 10% heat-inactivated fetal bovine serum, 100 U/ml penicillin and 100 μg/ml streptomycin, at 37°C in humidified atmosphere with 5% CO_2_. Cells were transfected 24h before further processing using the Amaxa Nucleofector® device (Lonza) following the manufacturer’s protocol.

### DNA Constructs

Constructs of HA-tagged murine NL1, NL2 and NL3 were a gift from Peter Scheiffele (Addgene plasmid #15260, #15246, and #59318, respectively (Chih et al., 2006, Chih et al., 2004)). All NL2 phosphorylation mutants (^HA^NL2-S714A, S714D, S714/719/721A, S714/719/721D, Y770A) were generated using this NL2 cDNA as a template, where subsequent mutagenesis was performed by DNAExpress (Canada). To create GST-NL2^CT^ WT and S714D, a PCR product of the intracellular domain flanked by *Bam*HI and *Xho*I restriction sites was created from the equivalent full-length construct (forward primer catcatggatcctacaagcgggaccggcgcc; reverse primer catcatctcgagctatacccgagtggtggagtg) and subcloned into pGEX4T3 (GE Healthcare). GFP-tagged Gephyrin-FingR was a gift from Don Arnold (Addgene plasmid #46296 (Gross et al., 2013)). eGFP-C1 used to identify transfected cells in electrophysiology experiments has been described previously (Hannan et al., 2020).

### Purification of GST-fusion protein and *in vitro* phosphorylation

GST fusion proteins were produced as described (Kittler et al., 2000). In brief, BL21 *E.Coli* containing pGEX4T3-NL2^CT^ WT or S714D were grown in Luria Broth until an OD600 of 0.5-0.6. Protein production was induced by addition of 1mM IPTG for 3h. Cells were harvested by centrifugation for 30 min at 4°C, washed in buffer containing 50 mM Tris pH8.0, 25% sucrose, 10 mM EDTA, and pelleted by centrifugation for 30 min at 4°C. Cells were then lysed by sonication in buffer containing 1% Triton-X100, 10 mM Tris pH7.4, 1 mM EDTA, 1 mM DTT, 1 mM PMSF, antipain, pepstatin and leupeptin at 10 μg/ml. Lysates were further incubated for 30 min upon addition of 12.5 mM HEPES pH 7.6, 75 mM KCl, 125 mM EDTA, 12.5% glycerol, and then spun for 1h at 12,000×*g* and 4°C. GST protein was purified by adding Sepharose 4B beads (GE Healthcare) and incubating 2h at 4°C. Beads were washed and stored at 4°C in buffer containing 20 mM HEPES pH7.6, 100 mM KCl, 0.2M EDTA, 20% glycerol, 1 mM DTT, and 1 mM PMSF. To perform *in vitro* phosphorylation 5 μg of purified GST-NL2^CT^ (WT or S714D) fusion protein was incubated with 2.5kUnits purified catalytic subunit of PKA (NEB) supplemented with 200 μM ATP at 30°C for the indicated time points.

### Protein sample preparation and western blotting

Neuronal and COS cell cultures were treated with DMSO (vehicle control, 1:1000 culture volume), 25 μM forskolin (Calbiochem), and/or 50 μM H89 (Sigma) for the time points indicated. Cell lysates were obtained by direct lysis in SDS sample buffer and boiling for 30 min prior to separation by gel electrophoresis. For *in vitro* dephosphorylation, untreated cultures of cortical neurons (DIV8) were lysed in lambda phosphatase buffer (NEB), incubated for 20 min to ensure total lysis, and spun for 10 min at 14,000×*g* to remove cell membranes. The supernatant was incubated with 500 Units lambda phosphatase (NEB) or phosphatase inhibitors (1 mM NaVO4, 5 mM NaF) for 1h at 30°C. The reaction was stopped by addition of SDS sample buffer.

For surface biotinylation, cortical cultures (DIV8) were treated for 1h with DMSO or forskolin. Biotinylation was performed as described (Arancibia-Carcamo et al., 2006). In brief, cells were washed 3x with ice-cold PBS supplemented with Mg^2+^ and Ca^2+^, then incubated with 0.5 mg/ml EZ-link Sulfo-NHS-LC-Biotin (ThermoFisher) for 12 mins on ice. The reaction was quenched using 1 mg/ml BSA in PBS, and then the cells were washed with PBS. Cells were lysed in RIPA buffer (150 mM NaCl, 1% Nonidet-P40, 0.5% DOC, 0.1% SDS, 1 mM EDTA, 2 mM EGTA, 1 mM PMSF, 50 mM Tris pH 7.5 supplemented with antipain, pepstatin and leupeptin at 10 μg/ml), and left to rotate at 4°C for 1h. Membranes were pelleted by centrifugation at 14,000×*g* for 10 minutes at 4°C. To isolate biotinylated surface proteins, supernatants were then incubated with 25 μl of a 50% Neutravidin bead slurry (Pierce, ThermoFisher) for 2h at 4°C. Beads were washed 3x in RIPA buffer before resuspending in sample buffer and analysed by SDS-PAGE and western blotting.

Protein samples were separated by standard Laemli SDS-PAGE on 9% Tris-Glycine gels and transferred onto nitrocellulose membrane (GE Healthcare Bio-Sciences). PhosTag gels were prepared by adding 200 μM PhosTag™ (WAKO chemicals, Alpha Laboratories) and 40 μM MnCl2 to a 5% standard Laemli SDS-PAGE gel. For detection by western blot, PhosTag gels were transferred to PVDF membrane (GE Healthcare). Membranes were blocked for 1h in milk (PBS, 0.1% Tween, 4% w/v skimmed milkpowder), and then incubated overnight at 4°C with shaking in primary antibodies diluted in milk (1:1000 Rabbit-anti-Neuroligin2 (Synaptic Systems 129-202), or 1:200 Mouse-anti-HA-tag (supernatant, clone 12CA5, produced in house). Blots were then incubated with the appropriate HRP-conjugated secondary antibody for 1h at room temperature, and developed using Luminate Crescendo Western HRP substrate (Millipore). Signal was detected using an ImageQuant LAS4000 mini (GE Life Sciences).

### Immunocytochemistry and Proximity Ligation Assay

Antibody feeding in hippocampal neurons (DIV13-14) was performed by adding 1:50 Mouse-anti-HA (supernatant, clone 12CA5, produced in house) directly to neuronal maintenance medium and allowing internalisation for 40 min at 37°C. Neurons were washed twice in medium before fixation for 7 min in 4% PFA (PBS, 4% paraformaldehdye, 4% sucrose, pH 7) and blocking for 10 min (PBS, 10% horse serum, 0.5% BSA). Surface ^HA^NL2 was stained for 45 min with 1:300 goat-anti-mouse Alexa555. Cells were then permeabilised by incubation in block solution supplemented with 0.2% Triton X-100 for 10 min, and subsequently incubated for 45 min with 1:500 donkey-anti-mouse Alexa488, and finally mounted using ProLong Gold antifade reagent (Invitrogen).

For cluster staining of surface-exposed NL2 and GABA_A_R γ2 subunit, hippocampal neurons (DIV13-14) were fixed and incubated in block solution as described above. Neurons were then stained for 1h at RT with primary antibody diluted in block solution (1:100 Rabbit-anti-NL2 (Alomone Labs, ANR-036) or 1:500 Guinea Pig-anti-γ2 (Synaptic Systems, 224-004)). Cells were then permeabilised in block solution supplemented with 0.2% Triton X-100, and incubated for 1h at RT with 1:1000 mouse anti-MAP2 (Synaptic Systems, 188-011), followed by 45-60 min incubation with the appropriate Alexa-fluorophore conjugated secondary antibodies (goat-anti-mouse Alexa405, goat-anti-rabbit Alexa555, and goat-anti-guinea pig Alexa647). Finally, coverslips were mounted onto glass slides using ProLong Gold antifade reagent (Invitrogen).

For Proximity Ligation Assays (PLA) hippocampal neurons (DIV13-14), where applicable treated with DMSO or forskolin for 1h, were fixed as described above, washed 3 times in PBS, and blocked for 10 mins in blocking buffer (10% Horse Serum, 5 mg/ml BSA, and 0.2% Triton in PBS). Coverslips were then incubated for 1h at room temperature with primary antibodies in blocking buffer (1:500 Rabbit anti-NL2 (Synaptic Systems, 129-203), or 1:400 Mouse anti-MPM2 (Millipore, 05-368), or 1:500 mouse anti-Gephyrin (Synaptic Systems, 147-011), or a combination of NL2 with MPM2 or NL2 with Gephyrin, in all cases combined with 1:1000 Guinea Pig anti-MAP2 (Synaptic Systems, 188-004). The remaining of the staining was performed according to the manufacturer’s protocol (Duolink® PLA). In brief, coverslips were incubated in a humidity chamber with anti-mouse MINUS and anti-rabbit PLUS probes (Sigma Aldrich) plus 1:500 goat anti-guinea pig Alexa647 for 1h at 37 °C, hybridized for 30 min at 37 °C, and amplified for 100 min at 37 °C using the red fluorophore (equivalent to TexasRed). Coverslips were then allowed to dry at room temperature for 20 min in the dark, and mounted using Duolink® *In Situ* mounting medium containing DAPI.

### Image acquisition and analysis

Confocal images were acquired on a Zeiss LSM700 upright confocal microscope using a 63x oil objective (NA: 1.4) and digitally captured using ZEN LSM software (version 2.3), with excitation at 405 nm for DAPI and Alexa-Fluor405, 488nm for GFP and Alexa-Fluor488, 555 nm for Alexa-Fluor555 and PLA red fluorophore (equivalent to TexasRed), and 633 nm for Alexa-Fluor647 conjugated secondary antibodies. Image analysis was performed using in-house written scripts for ImageJ (*https://imageJ.net/Welcome*), details of which are given below.

For antibody feeding, a single image with an optimal thickness of 1 μm was acquired for entire neurons (image size 203 × 203 μm) or a zoomed image of the soma (40.6 × 40.6 μm). Laser settings were adjusted to ensure channels were not saturated. Acquisition settings and laser power were kept constant within experiments. For quantification of antibody feeding, zoomed soma images of surface and internal staining were thresholded to generate a mask, which was then used to measure total intensity of surface and internal signal in the original image. The ratio between internal and total (surface plus internal) intensity was calculated as a relative measure for internalisation.

For PLA images, a single image with an optimal thickness of 1 μm was acquired for entire neurons (image size 201 × 201 μm). The field of view was selected based on MAP2 staining, ensuring each image contains a similar cell density and avoiding bias. Using ImageJ, a suitable threshold was selected for MAP2 and PLA signal and kept constant for all data sets. The number of PLA dots per MAP2 area was normalised to the average of the control condition per experiment (summed background in Fig.1e-f; DMSO-treated cells in Fig.3o-p).

For cluster analysis, 3 sections of dendrite per hippocampal neuron, ~50 μm from the soma, were imaged with a 3.4x zoom (equating to 30 μm length of dendrite). Cluster analysis was performed as described (Smith et al., 2014, Davenport et al., 2017, Halff et al., 2019). In brief, a suitable threshold was selected for each channel using Metamorph software (version 7.8; Molecular Devices) and applied to all images within the same dataset. Clusters with an intensity above this threshold and size of at least 0.04 μm^2^ were quantified. Quantification was performed on 5-8 cells per experiment.

### Single particle tracking

Single particle analysis was carried out by tagging ^HA^NL2 with quantum dot (QD) nanocrystals in primary hippocampal neurons (DIV13). Coverslips containing neurons expressing ^HA^NL2-WT or ^HA^NL2-S714A and GFP-Gephyrin-FingR were washed in Krebs solution (containing in mM: 140 NaCl, 4.7 KCl, 2.52 CaCl_2_, 1.2 MgCl_2_, 11 glucose, and 5 HEPES; pH 7.4) and subsequently incubated for 5 min at 37°C with 1:50 mouse anti-HA primary antibody (supernatant, clone 12CA5, produced in house) combined with DMSO (vehicle control, 1:1000 culture volume) or 25 μM forskolin in Krebs solution. Coverslips were then washed and incubated for 1 min at RT with 1:5000 anti-mouse secondary antibody conjugated via Streptavidin with Qdot655 (Invitrogen) in Krebs solution supplemented with 10% Horse Serum and 1:1000 DMSO or forskolin. After washes the coverslips were loaded in an environmental chamber (Solent Scientific) and imaged at 37°C within 5 min of mounting in a Krebs solution supplemented with DMSO or forskolin.

^HA^NL2-QD complexes were imaged using a wide-field inverted microscope (Olympus IX71) with a high magnification objective (60X; NA – 1.35; Olympus), a halogen light source (PhotoFluor-II Metal Halide illumination system) and a back-illuminated cooled electron-multiplying charge coupled device (EMCCD; iXon3 885, Andor Technology). GFP-Gephyrin-FingR was imaged with a 457–487 nm band-pass excitation filter (all filters / mirrors were acquired from Semrock), a 496 nm long-pass emission filter, and 495 nm dichroic mirror. QD655 was imaged with a 415-455 nm band-pass excitation filter, a 647.5-662.5 nm band-pass emission filter, and a 510 nm dichroic mirror. Real-time movement of NL2-QDs was captured by first imaging GFP-Gephyrin-FingR, followed by QD655s in the same region of interest and plane of view at 33 Hz for 300 frames.

Single particle analysis of ^HA^NL2-QDs was carried out as described previously (Hannan et al., 2016, Bannai et al., 2006). A custom Matlab (MathWorks) plugin, SPTrack (Ver5), was used to analyse QD movement by first identifying the centre of each QD using a 2-D Gaussian fit (spatial resolution ~10–20 nm). The centre of the Gaussian peaks of each QD in successive image frames were track-connected, based on estimated diffusion coefficients and the likelihood of the two Gaussian peaks in consecutive frames belonging to the same QD track. QDs trajectories with less than 15 consecutive frames were discarded.

The mean square displacement (MSD) of each QD was calculated using the following equation:

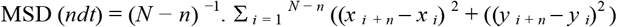

where *x*_*i*_ and *y*_*i*_ are the spatial co-ordinates of a single QD identified in a single image frame *i*. *N* is the total number of points in the QD trajectory, *dt* is the time interval between two successive frames (33 ms), and *ndt* is the time interval over which the mean square displacement is averaged. From the MSD plot, the diffusion coefficient, *D*, for a QD was calculated by fitting the first two to five points of the MSD plot against time with the following expression:

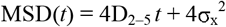

where σ_x_ is the QD localization accuracy in one dimension. D was determined from the slope of the relationship. Given the inherent noise in CCD imaging systems, and the errors in accurately locating single QDs that results, trajectories with D < 10^−4^ μm^2^ / s were considered immobile and excluded from calculation of median D.

Synaptic terminals, identified by GFP-Gephyrin-FingR puncta, were thresholded using ImageJ, and co-localised QD trajectories were defined as synaptic.

### Patch-clamp electrophysiology

Hippocampal neurons expressing ^HA^NL2 (WT or S714A or S714D) were identified by their co-expression with eGFP. Spontaneous inhibitory postsynaptic currents (sIPSCs) were recorded at DIV13–15 using patch clamp electrophysiology. Patch electrodes (4–5 MΩ) were filled with an internal solution containing (in mM): 120 CsCl 1 MgCl_2_, 11 EGTA, 30 KOH, 10 HEPES, 1 CaCl_2_, and 2 K_2_ATP; pH – 7.2. Neurons were superfused with a saline solution containing (mM): 140 NaCl, 4.7 KCl, 1.2 MgCl_2_, 2.52 CaCl_2_, 11 Glucose, and 5 HEPES; pH 7.4 supplemented with 2 mM kynurenic acid to block all spontaneous excitatory post-synaptic currents. Membrane currents were filtered at 5 kHz (−3 dB, 6th pole Bessel, 36 dB/octave) with optimised series resistance (Rs, <10 MΩ) and whole-cell membrane capacitance compensation. Membrane currents were recorded at a holding potential of −60 mV. Changes of series resistance greater than 10% during the experiment resulted in the recording being excluded from analysis.

sIPSCs were analysed using WinEDR and WinWCP (John Dempster, University of Strathclyde, UK). Amplitude distributions were fitted with a sum of 1–4 Gaussian functions according to the function:

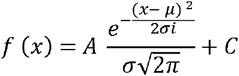

Where A is the amplitude and C defines the pedestal of the histogram. This function provided the Gaussian mean amplitude current (μ) and standard deviation (σ). All the distributions were fitted using this function in Origin (Ver 6). The accuracy of the fits was checked by repeating the iterative non-linear fitting procedure after substituting the best-fit.

### Statistical analysis

For all quantified fixed imaging experiments the experimenters were blind to the condition of the sample analysed. All quantified experiments were performed 3 times from independent cell preparations, treatments, and transfections, unless indicated otherwise in the figure legends. N-numbers indicate number of experiments (western blots, PLA for NL2+MPM2), number of trajectories (QD experiments) or number of cells (all other cases). Values are given as mean ± standard error of the mean (SEM) or median with Inter-quartile range (25-75%) as indicated in the figure legends. Error bars represent SEM.

Statistical analysis was performed in GraphPad Prism (version 8; GraphPad Software, CA, USA) or Microsoft Excel. All data was tested for normal distribution with D’Agostino & Pearson test to determine the use of parametric (student’s *t*-test, one-way ANOVA) or non-parametric (Mann-Whitney, Kruskal-Wallis) tests. When p<0.05, appropriate *post-hoc* tests were carried out in analyses with multiple comparisons and are stated in the figure legends.

## Data availability statement

The datasets generated and/or analysed during the current study are available from the corresponding author on reasonable request.

